# A new molecular method for the exploration of hybrid zones between two toad species of conservation interest

**DOI:** 10.1101/2020.07.29.226472

**Authors:** Zoltán Gál, Tibor Kovács, János Ujszegi, Brandon P. Anthony, Balázs Vági, Orsolya I. Hoffmann

## Abstract

Analyzing hybrid zones between previously isolated lineages allows insight into processes determining the fate of re-encounter of these taxa. The distributions of Fire-bellied (*Bombina bombina*) and Yellow-bellied toads (*B. variegata*) meet in the Carpathian Basin resulting in a narrow contact zone in the foothill regions, where hybrids often appear. Our aim was to explore a transient zone between *B. bombina* and *B. variegata* within the Carpathian Basin along a transect including the Börzsöny Hills in Hungary and Krupinská Planina in Slovakia. We visited 28 locations in these areas and collected altogether 230 specimens, photographed and sampled them using buccal swabs. In order to distinguish between the two species and determine hybrid individuals, we used mitochondrial markers and designed a novel technique based on the restriction of the Ncx-1 gene. The geographical distribution pattern of these two species delivered unexpected results, as Börzsöny Hills was completely colonised by *B. bombina* including locations which can be classified as typical habitats of *B. variegata*. Moreover, in Krupinská Planina many locations were also colonised by *B. bombina*, including high altitude ponds. The most remote sites still harbour *B. variegata* populations, but seven of these were found with hybrid individuals. This pattern may indicate the northward and altitudinal range expansion of *B. bombina* and the colonisation of habitats of its sister species in these areas. Our results warrant enhanced attention to hybrid zones, where introgression and changes in population composition may reflect recent rapid environmental alterations and redirect conservation focus.

## Introduction

One of the main processes driving speciation is geographic isolation due to one or more vicariant events leading to divergent lineages [1]. On one hand, different ecological constrains distinctly shape the two lineages, facilitating ecological differentiation (i.e. adaptation to different habitats). On the other hand, allopatric distribution may lead to the accumulation of random genetic differences between the isolated lineages. Both types of genetic divergence may potentiate reproductive isolation. However, if there is insufficient time available for speciation, incomplete reproductive isolation may result in hybridization between the sister-lineages upon re-encounter [2,3]. In Europe, several cases of hybridization are considered to be a consequence of warming periods after glacial ages, when previously allopatric populations of sister lineages became parapatric due to area-expansions and came into secondary contact [4]. Analyzing hybrid zones of genetically and ecologically differentiated, but not reproductively isolated taxa allows insight into admixture processes determining the fate of the closely related lineages (see e.g. [5]).

The hybrid zone between the European fire-bellied and yellow-bellied toads (*Bombina bombina* and *B. variegata*; Bombinatoridae) in Central Europe has been identified by morphologic, electrophoretic and genetic analyses [6–9]. The two species are similar with a warty, cryptically coloured dorsal and aposematic, brightly coloured ventral pattern [10,11]. The first attempts to distinguish between sister species and hybrids were based on morphological characteristics. The most commonly used trait was the extension and connectivity of the ventral patches at particular body parts, supplemented with morphometric measurements, including snout-vent length, head length, head width, distance between eyes, forelimb length, femur length, tibia length, and foot length [8,12]. However, transitional morphological characters between *B. bombina* and *B. variegata* had been observed as early as the 1890s [13], due to hybridization in the contact zones [9,14]. Therefore, more reliable identification based on molecular methods were designated including protein electrophoresis [7,9,15,16] and DNA techniques using mitochondrial [8,17] and nuclear gene markers [18] or microsatellites [19,20]. Molecular analyses revealed that introgression has not been detected outside the hybrid zones and the genetic structures vary in pure and hybrid populations [10,17–19,21,22]. The hybrid zones cover different genetic structures depending on their location [18].

The location of the hybrid zone is determined by environmental factors due to the diverse adaptations to different habitats of the sister species. The distribution of *B. bombina* covers the lowlands of Europe from Germany to Russia and probably extends even beyond the Ural Mountains, while *B. variegata* occurs mainly in the highlands of the Balkan Peninsula, Central and Western Europe and is absent east of the Carpathian Ridges [23]. Thus, the two species have parapatric distribution with a narrow but extended contact zone stretching from Germany southward to Bulgaria [24]. Based on mtDNA analyses, *B. bombina* is composed of two clades confined to Northern and Southern Europe, with low mtDNA divergence between and within clades [17,18]. In contrast, mtDNA variation of *B. variegata* in its range is extensive, and four distinct haplotypes are identified. Hence the overall pattern of their recent distribution reflects a post-glacial radiation from Balkan, Mediterranean and Carpathian refugia, whereas recolonization of European habitats by *B. bombina* initiated from one refugium near the Black Sea coast [17,18,25].

The Carpathians and their surrounded basin with complex topography created a complicated and diverse contact zone between the two species. The inner part of the Carpathian Basin is divided by mid-sized ridges with the highest elevation of Kékes (Hungary) up to 1015 m above sea level (asl), where *B. variegata* occurs in isolated populations inhabiting small temporary ponds and puddles at these mountainous enclaves. In contrast, previous studies indicate that *B. bombina* prefers larger, more permanent water bodies at lowland, open areas with more connected populations [21,26]. Due to the different habitat preferences, the two species contact mostly along the foothill regions, forming an extended contact zone in the Carpathian Basin with resulting hybrid individuals in many populations [8,16]. Gollmann and colleagues [7] studied a shorter section of the ridges on the border of northern Hungary and southern Slovakia in the Aggtelek Karst region and revealed a hybrid zone between the two species with highly variable population structures. Sas and colleagues [11] reported a pure hybrid population around Oradea, Romania. Evidence of hybridizing populations from other sections of the Carpathians were also shown [17,19,22].

Our aim was to explore a transient zone between *B. bombina* and *B. variegata* and assess the distribution of the two species and their hybrids from the inner part of the Carpathian Basin to the southern edge of the Carpathian Ridges along an approximately 50 km long north-south transect (Fig 1). We also aimed to test a novel molecular approach to distinguish between the two species and determine hybrid individuals.

**Fig 1.**
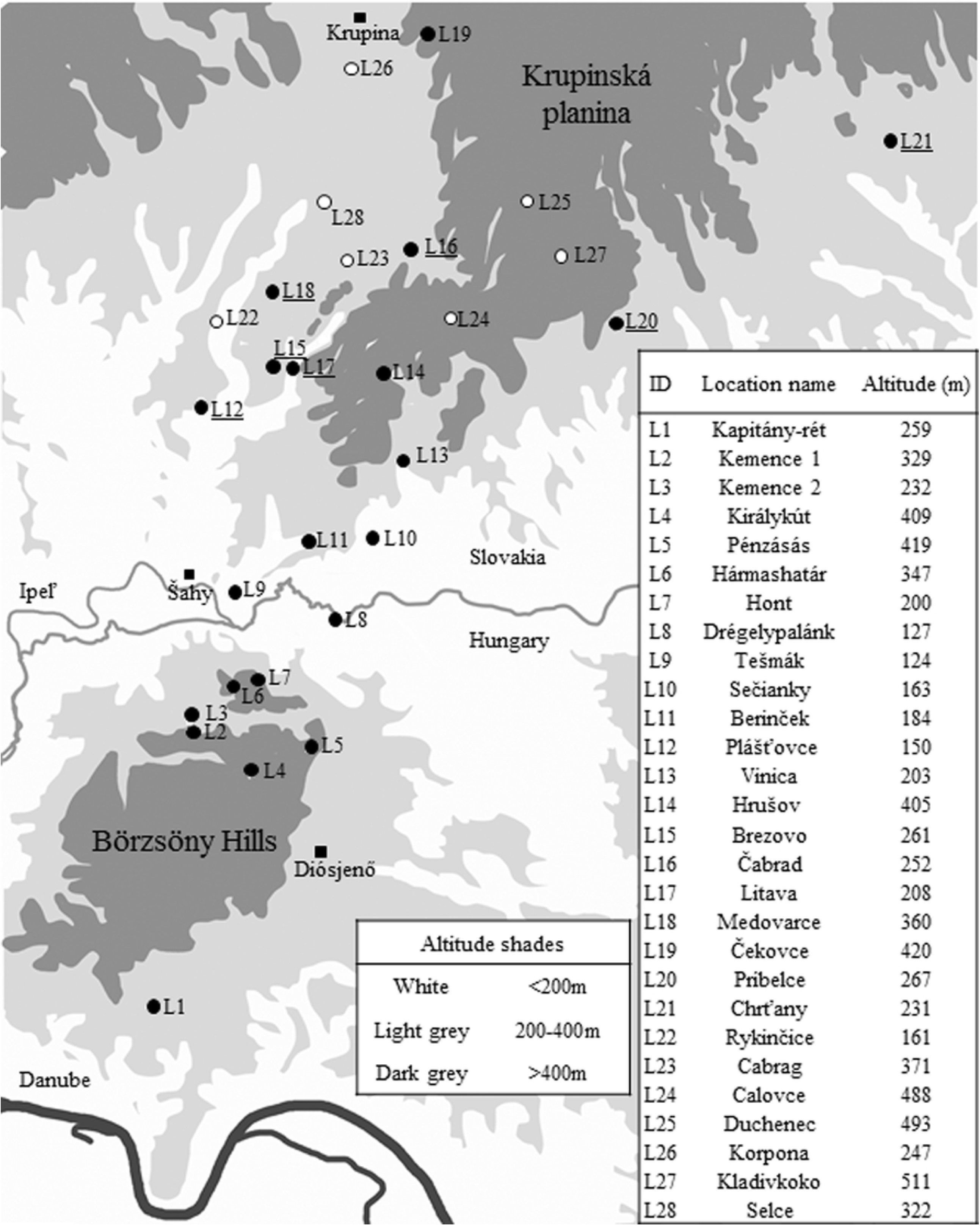
Map of the sampling area. Circles indicate the assessed locations (L1-21) and empty circles represent locations where individuals were not observed (L22-L28). Squares represent settlements. White, light grey and dark grey areas represent the relief under 200 m, between 200-400 m and above 400 m, respectively. Locations containing hybrid individuals are underlined.

## Materials and Methods

### Study area

Börzsöny Hills are located in northern Hungary. In the north they are separated from Krupinská Planina in Slovakia by Ipel’ River’s valley, which also demarcates the border between the two countries (Fig 1.). The highest elevation in Börzsöny is 938 m asl and, due to its rugged terrain, most of the valleys are strongly shaded with cold microclimates. Krupinská Planina is a sibling mountain of Börzsöny in Slovakia having identical rock material; a Miocene andesite. It is lower (535 m asl), but its terrain is heavily rugged as well. Published herpetofaunistical data from Börzsöny is old [27] and while plenty of data were collected in an online database (https://herpterkep.mme.hu), no systematic assessment was made for *Bombina* species prior to this study. The herpetofauna of Krupinská Planina had hardly been explored before we started our investigation. The only record was available from Slovakian sources is a summary of a 2-year (2008-2009) data collection. Both Bombina species had been found in the survey, but the author did not analyse their presence deeper [28].

### Sample collection

We searched for *Bombina* habitats based on literature sources and an online data base (herpterkep.mme.hu). Potential habitats were marked on satellite maps (GoogleEarth). In total, we visited 28 locations (S1 Table) and sampled 230 toads between 2014 and 2018. Individuals were caught by hand and DNA samples were collected using a non-invasive buccal swabbing. Each individual was placed into a transparent plastic box and fixed by gently pressing a plastic sponge against their back. The ventral pattern of each toad was photographed for further analyses (not presented here, S1 Fig and they were then released at the capture site).

### Amplification and analysis of the nuclear Ncx-1 gene

The collected swabs were stored in 70% ethanol until the DNA extraction by phenol-chloroform method [29]. The following primers were used to amplify a 846 base pair (bp) long region of the *Ncx-1* gene: NcxF (5’-TCATCCGCTCCTGAAATTCT -3’) and NcxR (5’-CACAGTCCCACAGTTTTCCA -3’) [18]. All PCR reactions were implemented with MyTaq Ready Mix (Bioline) according to the manufacturer’s instructions. The reactions contained 80 ng DNA and 8 μM of each primer. The PCR conditions were the following: an initial denaturation step at 95 °C for 5 min, 30 cycles with denaturation at 95 °C for 20 sec, an annealing temperature of 62 °C for 20 sec, and elongation at 72 °C for 20 sec and a final elongation at 72 °C for 2 min. The 846 bp long PCR product was analysed in 1% agarose gel.

To find a restriction fragment length polymorphism (RFLP) loci *Ncx-1* gene, haplotypes were downloaded from the National Center for Biotechnology Information (NCBI) and aligned to a consensus sequence both in *B. bombina* and *B. variegata* with ClustalW2 (EMBL-EBI). Three haplotypes were obtained from *B. bombina* and 16 sequences originated from *B. variegata* according to Fijarczyk et al. [18]. The GenBank accession numbers can be found in S2 Table. The created *B. bombina* and *B. variegata Ncx-1* consensus sequences were compared pairwise with MultAlign [30]. With NEBCutter [31] and dCAPS Finder [32] four HpyCH4V restriction enzyme sites (TG_^CA) were detected in *B. bombina*, with one out of four missing from the *B. variegata* sequences (S2 Fig). The amplified PCR fragments were digested with HpyCH4V enzyme (New England Biolab) at 37 °C for 3 hours. 10 μl of each digestion reaction were loaded into 2.5 % TAE agarose gel and GeneRuler™ 1 kb Plus DNA Ladder (Fermentas) was used as a molecular weight marker. The digestion results in a 694 bp long fragment in case of *B. variegata. B. bombina* samples cleave to a 369 bp and 325 bp long fragments. The samples of hybrid individuals carry both the 694 bp, 369 bp and the 325 bp long fragments. The remaining 152 bp split into 96 bp, 32 bp and 24 bp long fragments in both genotypes (Fig 2).

**Fig 2.**
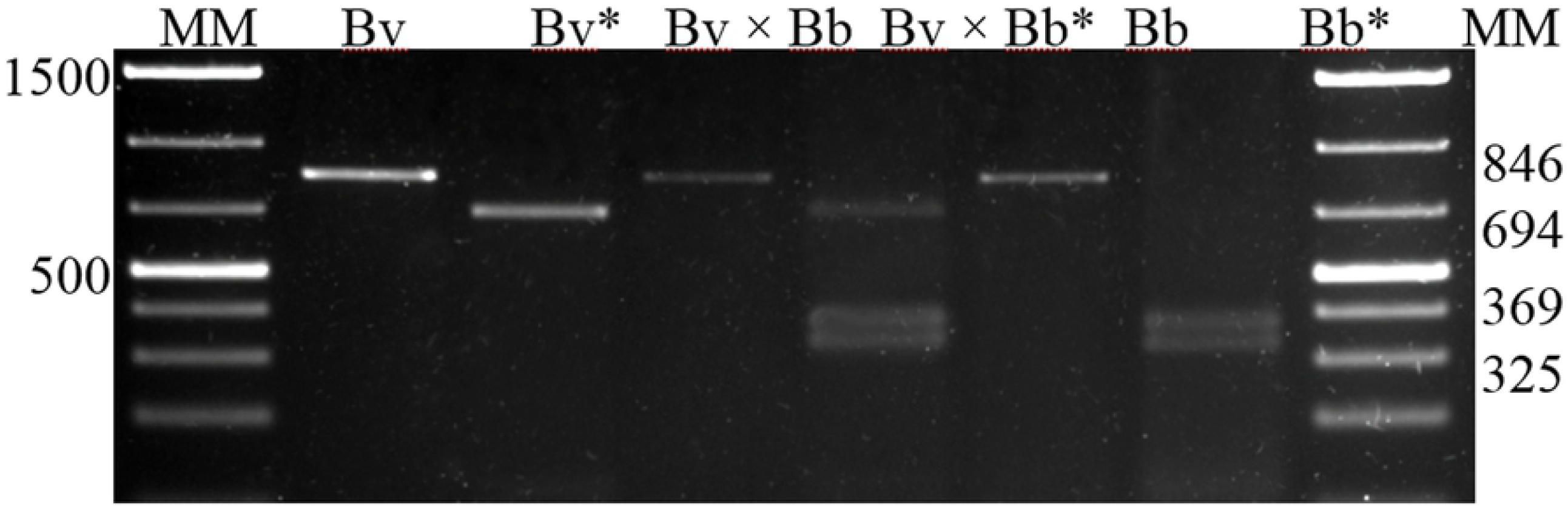
Electrophoretic differentiation of the digested PCR fragments specific to Ncx-1 gene. The amplified PCR product from Ncx-1 gene before enzymatic digestion: *B. variegata* (Bv; identifier of the individual: 101), *B. bombina* (Bb; identifier of the individual: 189) and hybrid (Bv × Bb; identifier of the individual: 169) Fragments resulted by the digestion with HpyCH4V restriction enzyme: Bv* (101), Bv × Bb* (169), Bb* (189). MM – molecular weight marker.

### Amplification of the mitochondrial fragments

Six samples were processed at each location where one species emerged. All the samples were analysed at the hybrid locations to detect the types of the mitochondria that occur. The primers were based on the total *B. bombina* and *B. variegata* mitochondrial genome at NCBI (EU115993.1, NC_009258.1). Two mitochondrial primer pairs were used to distinguish the two species: RadF (5’-CAGCTAGTATCAACCCACCAGAT – 3’) and RadR (5’-TTGATCTGTTGCTGGGTACGTCTTG -3’) primers are specific for *B. bombina*, TynF (5’-CAATAAAATTCAACCGCCAACAAT -3’) and TynR (5’-AAGTTGATCTGTTGCTGGGTATGTTCTA -3’) primers are specific for *B. variegata* mitochondria [33]. The PCR reactions were completed with Mytaq Read Mix (Bioline) according to the manufacturer’s instruction. The PCR cycles were the following in case of the Rad primer pair: an initial denaturation step at 95 °C for 5 min, 35 cycles with denaturation at 95 °C for 20 sec, an annealing temperature of 63 °C for 20 sec, and elongation at 72 °C for 20 sec and a final elongation at 72 °C for 2 min. With the Tyn primer pair the same PCR protocol was used except that the annealing temperature was 60 °C. The resulting DNA fragments were analyzed at 2.5 % agarose gel containing ethidium-bromide with GeneRuler™ 1 kb Plus DNA Ladder (Fermentas). The RadF-R primers amplify a 173 bp long fragment, the amplicon with the TynF-R primers is 196 bp long (Fig 3).

**Fig 3.**
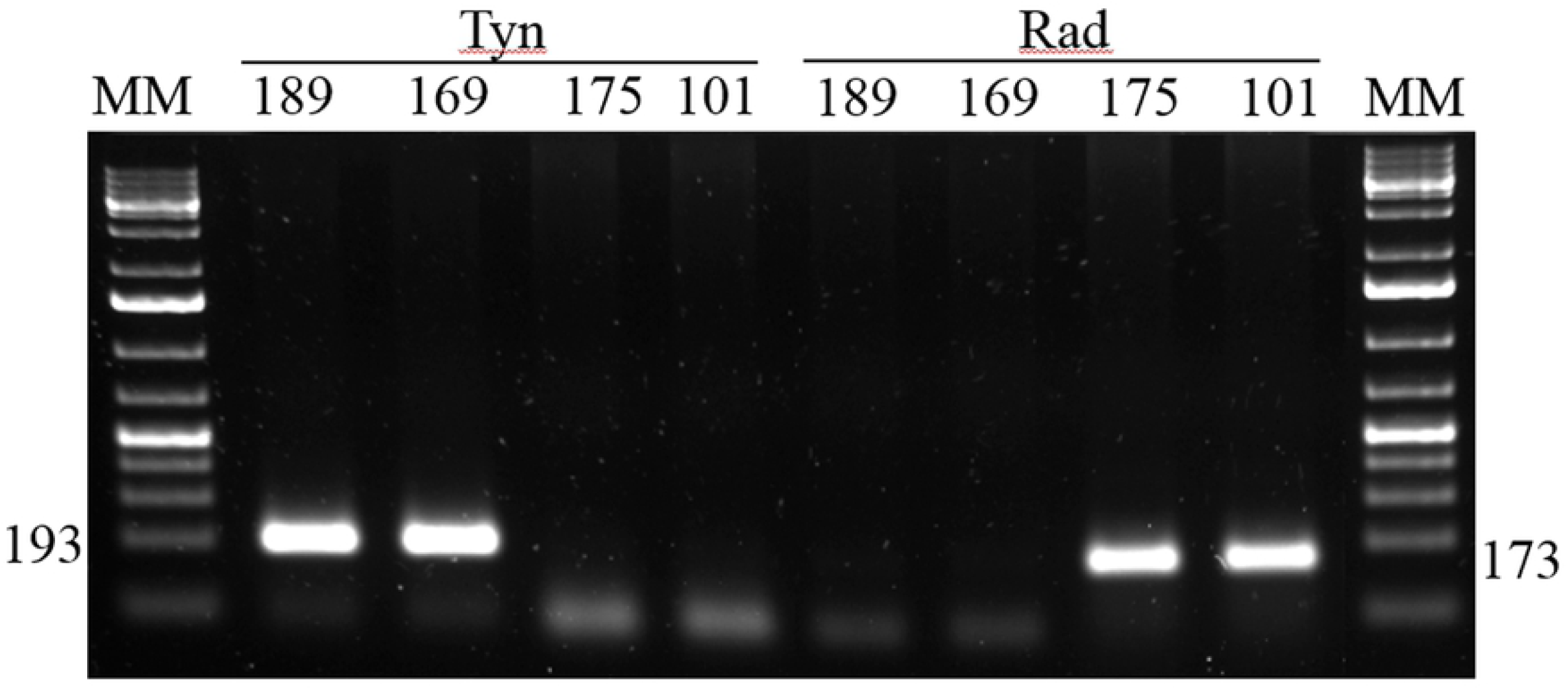
The amplification of mitochondrial regions with Tyn and Rad primer pairs specific for *B. bombina* (Bb) and *B. variegata* (Bv) respectively. According to the mithocondrial marker the individual 189 carries Bb genotype, while the individual 101 carries Bv genotype. Individual 169 and 175 are hybrids. MM – molecular weight marker.

### Sequencing

The PCR amplified *Ncx-1* gene fragments were sequenced by Sanger sequencing (Eurofins Genomics, Ebersberg, Germany), and resulting chromatograms analysed with Chromas 2.6.5 software (Technelyium).

## Results

### Identifying a new nuclear marker in Ncx-1 gene

Altogether, we analysed 230 samples from 21 sampling sites located in Hungary and Slovakia (Table 1). We could successfully amplify an 846 bp long fragment of the *Ncx-1* gene in all 230 cases. The designed restriction enzyme digestion with HpyCH4V could distinguish between the consensus sequences assigned to the two species and the hybrid individuals. To validate our results, we Sanger sequenced 4 *B. variegata*, 7 *B. bombina* and all the 26 hybrid individuals. The sequencing confirmed the result of the *Ncx-1* digestion in all cases as the heterozygous locus was specific to the individuals which were considered hybrids based on the digestion fragment composition (S2 Fig).

**Table 1.** Species identification results according to nuclear marker Ncx-1 at each location.

| Location | Sample size | Nuclear marker |  |  |
| --- | --- | --- | --- | --- |
|  |  | Bb | Bv | H |
| L1 | 29 | 29 | - | - |
| L2 | 1 | 1 | - | - |
| L3 | 2 | 2 | - | - |
| L4 | 11 | 11 | - | - |
| L5 | 15 | 15 | - | - |
| L6 | 5 | 5 | - | - |
| L7 | 10 | 10 | - | - |
| L8 | 10 | 10 | - | - |
| L9 | 15 | 15 | - | - |
| L10 | 5 | 5 | - | - |
| L11 | 6 | 6 | - | - |
| L12 | 19 | 14 | - | 5 |
| L13 | 20 | 20 | - | - |
| L14 | 15 | 15 | - | - |
| L15 | 3 | 2 | - | 1 |
| L16 | 9 | - | 6 | 3 |
| L17 | 34 | 6 | 14 | 14 |
| L18 | 7 | - | 6 | 1 |
| L19 | 2 | - | 2 | - |
| L20 | 10 | 5 | 4 | 1 |
| L21 | 2 | - | 1 | 1 |
| **Sum** | **230** | **171** | **33** | **26** |

Abbreviations: Bb: *Bombina bombina*, Bv: *Bombina variegata*, H: hybrid individual

### mtDNA markers

One hundred and nineteen samples were analysed with two mitochondrial markers from 14 locations (including all 7 locations with hybrids and half of the locations where only *B. bombina* was found) with the two primer pairs that can distinguish between the *B. bombina* and *B. variegata* mitochondria. The analysed 119 samples showed that the phenotype and the *Ncx-1* gene results agreed with the mitochondrial markers at all 7 tested locations where we found only *B. bombina* individuals (Table 1). Among the 7 sites with hybrids, at one location the mitochondrial markers revealed *B. bombina* genotype, at five locations the results showed *B. variegata* mitochondria. There was one location at the hybrid zone – Plášt’ovce (L12) – where we detected both *B. bombina* and *B. variegata* mitochondrion. Table 2 summarizes the result of the analyses based on the two mitochondrial markers in 22 hybrid individuals (in 4 individuals we failed to amplify mtDNA; see S3 Table for more details).

**Table 2.** Combined mitochondrial and nuclear genotypes of individuals from locations where hybrids occured.

| Nuclear marker | Mitochondrial marker | Sample size | Location ID |
| --- | --- | --- | --- |
| Bb | Bb | 14 | L12, L15, L17 |
| Bb | Bv | 7 | L17, L20 |
| Bv | Bv | 27 | L16, L17, L18, L20, L21 |
| H | Bb | 3 | L12, L15 |
| H | Bv | 19 | L12, L16, L17, L18, L20 |

Abbreviations: Bb: *Bombina bombina*, Bv: *Bombina variegata*, H: hybrid individual

### Distribution of the two species in the study area

Based on our species identification method, *Bombina bombina* occurred at 17 locations (including all locations in Börzsöny Hills), while *B. variegata* occurred at six locations. In 13 locations all individuals proved to be *B. bombina*. At one location (Čekovce) we found only *B. variegata*. We identified 7 locations where hybrid individuals occurred. In total, we caught 26 hybrid individuals identified with the *Ncx-1* nuclear gene marker at seven locations: Plášt’ovce, Brezovo, Čabrad, Litava, Medovarce, Pribelce, Chrtany. At Litava and Príbelce we found both *B. bombina, B. variegata* and also hybrid toads.

The examined toad populations in the Börzsöny Hills and the Ipel’ Valley consist of *B. bombina* specimens, and both species were found in the Krupinská Planina area. *B. variegata* appears exclusively in the coolest, remote mountainous areas of the region, while the edges of the plateau at even higher elevations were occupied by *B. bombina*. On average, *B. variegata* occupied higher elevations than *B. bombina*, but the two species reached the same maximum altitude. Hybrids occurred at intermediate altitude between the two species median elevations (Fig 4). However, there was no significant difference between the altitudinal distributions of the two species and their hybrids (one-way ANOVA, for all location: *F*_2,27_ = 0.473; P = 0.63; only for locations in Krupinská Planina, exluding *B. bombina* sites in Börzsöny and Ipel’ Valley: *F*_2,18_ = 1.064; P = 0.37). In the northern half of the Krupinská Planina we investigated 7 more locations (all artificial ponds), where we did not find any *Bombina* specimens (Fig 1).

**Fig 4.**
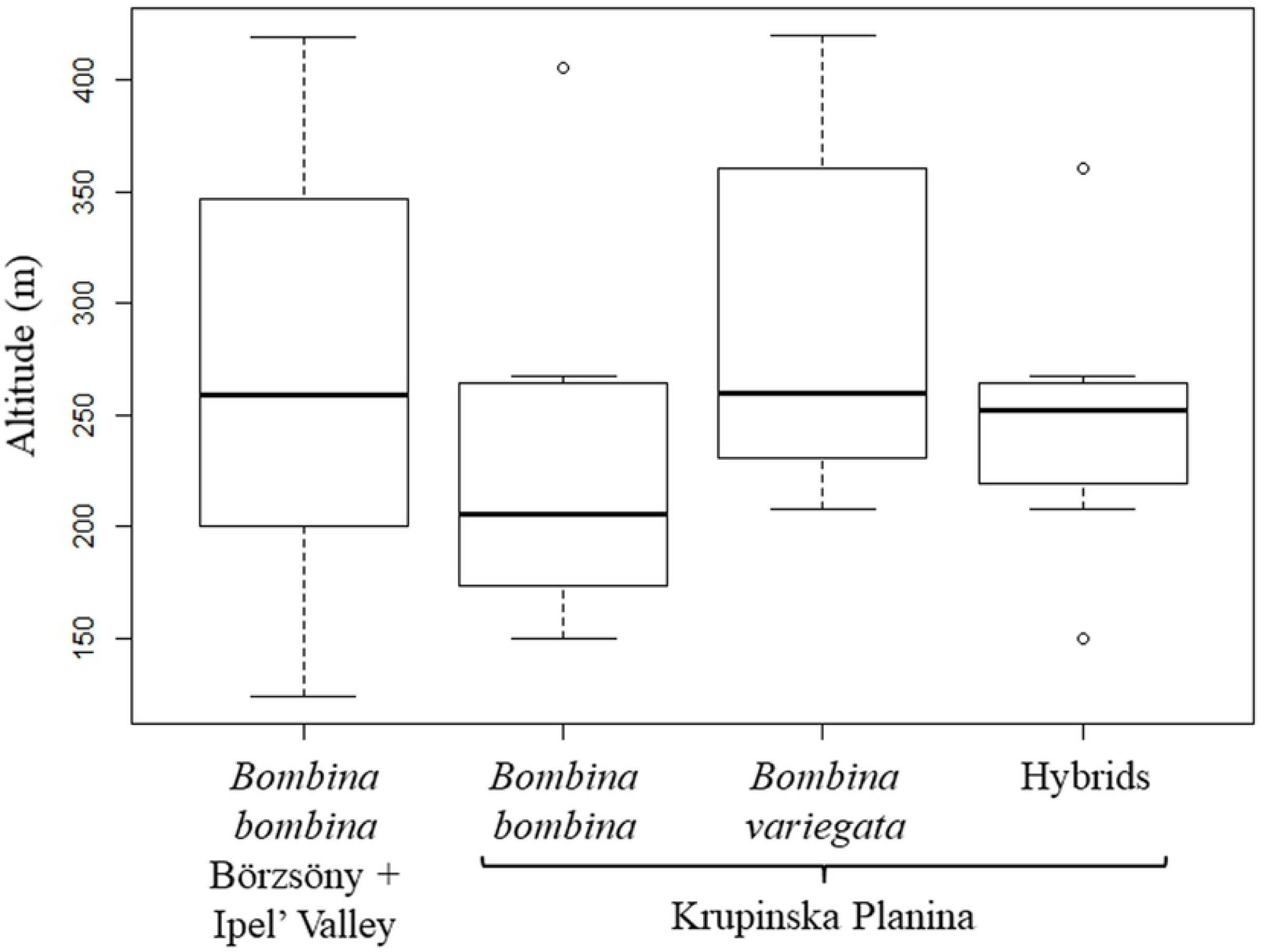
The altitudinal distribution of the two species and their hybrids in the two areas. Horizontal line is the median, whiskers represent range, boxes represent interquartiles and the circle indicates an outlier (deviating from the boundary of the interquartile range (IQR) by more than 1.5×IQR).

## Discussion

Our study has four main achievements. First, using restricted digestion of a nuclear gene we invented a new genetic test for the identification of *B. bombina* and *B. variegata* and their hybrids. Second, we successfully applied this method on *Bombina* samples from a putative hybrid zone and identified the two homozygous and the heterozygous genotypes. Third, we compared the nuclear genotypes of the sampled individuals with their haplotypes to a mitochondrial marker and found various genomic combinations in the actual hybrid zone. Fourth, we identified that, in contrast to Börzsöny Hills, Krupinská Planina is inhabited by both *B. bombina* and *B. variegata* populations which produce hybrids in their contact zones.

### Evaluation of molecular identification

The distribution patterns we explored are characteristically different at the two sides of the Ipel’ Valley. While in Börzsöny Hills (Hungary) we found only pure *B. bombina* populations even at the highest locations and in the smallest wheel track water bodies, in Krupinská Planina (Slovakia) the distribution of the two species and hybrids show a clear geographic and altitudinal pattern. *B. bombina* occupies the lowest and more southern locations, while *B. variegata* occurs at the most northern location and highest altitude, while intermediate locations (both in altitudinal and geographic terms) are inhabited by hybrid populations, where *B. variegata* appear to be the more abundant parent species. It is an interesting question if Börzsöny Hills harboured *B. variegata* populations in the past. Today’s remaining *B. variegata* populations in northern Hungary are mostly found in ranges including Zemplén or Aggtelek which are continuous with the Slovakian Carpathians. However, there is still *B. variegata* in Mátra Hills, a range very similar to Börzsöny in its more isolated location and geography. If *B. variegata* occurred in Börzsöny Hills in the past, it has been most likely outcompeted by its congener, which may be a potential genetic outcome of the introgression if environmental conditions favour one of the species. However, we did not detect hybridization introgression, as we only found pure *B. bombina* specimens in Börzsöny, based on both the nuclear and mitochondrial genomes.

In contrast, in the Krupinská Planina the greatest frequency of hybrid individuals has *B. variegata* mitochondrial haplotype. This suggests that most hybrids have *B. variegata* maternal origins, implying that hybridization arose from matings between *B. variegata* females and *B. bombina* males. This is consistent with the pattern in other frog species, where males disperse further than females [34], and that indiscriminate, coercive male frogs readily mate with heterospecific females both under experimental and natural conditions [35,36]. As hybrids occurred mostly alongside *B. variegata* individuals, it seems likely that *B. bombina* genes introgressed into these populations via occasional dispersal of *B. bombina* individuals.

Climate change will alter (or most probably already is altering) the recent pattern of habitats and biotopes of Europe and the entire globe as it has become clear by the end of the 20^th^ century [37,38]. As a response, several species will certainly shift their range adapting to the new circumstances [39,40]. Although in many cases studies revealed rapid and sometimes irreversible decline in amphibian populations as a consequence of global warming [41–43], some species will respond by dispersing and expanding their ranges [44]. The current isolated distribution patches of *B. variegata* probably resulted from the recent spreading of its congener, which outcompeted it from lower elevations of its former distribution area [8]. This scenario may also have occurred in the Börzsöny Hills, according to our results. In our case, we posit that if characteristically lowland species (*B. bombina*) colonize habitats where the ecological conditions would be more favourable for highland species (*B. variegata*), it reflects a current area expansion of the first species. Possible area expansion of *B. bombina* has been documented during the last decade at the northern edge of its range [45]. In Brandenburg province of Eastern Germany a detailed analysis suggested a possible area expansion of *B. bombina*, as further habitats are becoming suitable for the species due to predicted changes in climate and associated future environmental conditions [46].

As we did not investigate *Bombina* dispersal processes between ponds in our study area, nor their individual or environmental factors, we can only speculate that absence of either species in the artificial ponds in the northernmost portion of our study site is a result of (i) the relatively short time since these ponds have been established coupled with relatively short dispersal distances of *Bombina* species [47], (ii) barriers to dispersal within the landscape matrix including unsuitable microclimates and habitat for facilitating movement [48], and/or (iii) the fact that these permanent ponds were artificial, deeper and moderate in size which are less likely to be utilized by *B. variegata* in such contexts [49]. Nevertheless, we feel that extended investigation of ongoing patterns of dispersal, occupancy and interspecific dynamics in such hybrid zones is warranted.

Both *Bombina* species are considered reliable indicators of habitat quality and listed under the European Union’s Council Directive 92/43/EEC on the conservation of natural habitats and of wild fauna and flora (https://eur-lex.europa.eu). Being species of community interest, their population sizes should be regularly monitored. Our research highlights that special attention should be paid to hybrid zones, where changes of population composition may reflect effects of recent rapid environmental alterations, most notably climatic changes. As morphological identification of individuals could be problematic due to the large overlap between phenotypes, our new and simple genetic method provides a useful tool to track the hybrid populations genetic composition.

## Acknowledgements

Mitochondrial primers were kindly provided by the Jacek M. Szymura lab. The authors would like to thank to Peter Mikulicek and Daniel Jablonski for their advices on *Bombina* habitats in Slovakia and their help in receiving the Slovak licence, Attila László Péntek, Anna Fekete, Marietta Tóth, Márta Egyed and Tímea Bozsoky for their help during data collection.

## Supporting information

**S1 Fig. Photos of three individuals (Bv, Bv x Bb, Bb).** Sampling locations and individual IDs are indicated at the photos.

**S2 Fig. Side-by-side alignment of *Bv* consensus and *Bb* consensus sequence of the *Ncx-1* gene.** Stars represent the distinct nucleotides. Underlined regions mark the recognition site of the HpyCH4V restriction enzyme.

**S1 Table. The name and geocoordinates of sampling locations.**

**S2 Table. GenBank accession numbers of the sequences used to align the consensus sequence of the *Ncx-1* gene.**

**S3 Table. Combined mitochondrial and nuclear genotypes of individuals at each locations where hybrids were found.** In case of 2 individual in L12, 6 individuals in L20 and 1 individual in L17, L18 and L21 we failed to amplify any mtDNA. In L12, L17 and L18 the individuals was 1hybrids according to the nuclear marker.

**S4 Table. Sample size of individuals of which mtDNA amplification was failed at each location of the hybrid zone**

## References

[1] Riddle BR, Hafner DJ, Alexander LF, Jaeger JR. Cryptic vicariance in the historical assembly of a Baja California peninsular desert biota. Proc Natl Acad Sci U S A 2000;97:14438–43. https://doi.org/10.1073/pnas.250413397.

[2] Barton NH, Hewitt GM. Analysis of Hybrid Zones. Annu Rev Ecol Syst 1985;16:113–48. https://doi.org/10.1146/annurev.es.16.110185.000553.

[3] Harrison RG. Hybrid zones and the evolutionary process. Oxford Univ Press 1993:364.

[4] Hewitt G. The genetic legacy of the Quaternary ice ages. Nature 2000;405:907–13. https://doi.org/10.1038/35016000.

[5] Arntzen JW, Canestrelli D, Martínez-Solano I. Environmental correlates of the European common toad hybrid zone. Contrib to Zool 2020;89:270–81. https://doi.org/10.1163/18759866-bja10001.

[6] Szymura JM, Barton NH. Genetic analysis of a hybrid zone between the Fire-bellied toads, *Bombina bombina* and *B. variegata*, near Cracow in Southern Poland. Evolution (N Y) 1986;40:1141–59. https://doi.org/10.1111/j.1558-5646.1986.tb05740.x.

[7] Gollmann G, Roth P, Hodl W. Hybridization between the fire-bellied toads *Bombina bombina* and *Bombina variegata* in the karst regions of Slovakia and Hungary: morphological and allozyme evidence. J Evol Biol 1988;1:3–14. https://doi.org/10.1046/j.1420-9101.1988.1010003.x.

[8] Vörös J, Alcobendas M, Martínez-Solano I, García-París M. Evolution of *Bombina bombina* and *Bombina variegata* (Anura: Discoglossidae) in the Carpathian Basin: A history of repeated mt-DNA introgression across species. Mol Phylogenet Evol 2006;38:705–18. https://doi.org/10.1016/j.ympev.2005.08.010.

[9] Szymura JM. Hybridization between Discoglossid toads *Bombina bombina* and *Bombina variegata* in southern Poland as revealed by the electrophoretic technique. J Zool Syst Evol Res 2009;14:227–36. https://doi.org/10.1111/j.1439-0469.1976.tb00938.x.

[10] Szymura JM, Barton NH. The genetic structure of the hybrid zone between the fire bellied toads *Bombina bombina* and *B. variegata* : comparison between transects and between loci. Evolution (N Y) 1991;45:237–61. https://doi.org/10.1111/j.1558-5646.1991.tb04400.x.

[11] Sas I, Covaciu-Marcov S-D, Pop M, Ile R-D, Szeibel N, Duma C. About a closed hybrid population between *Bombina bombina* and *Bombina variegata* from Oradea (Bihor county, Romania). North West J Zool 2005;1:41–60.

[12] Gollmann G. Allozymic and morphological variation in the hybrid zone between *Bombina bombina* and *Bombina variegata* (Anura, Discoglossidae) in northeastern Austria. J Zool Syst Evol Res 2009;22:51–64. https://doi.org/10.1111/j.1439-0469.1984.tb00562.x.

[13] Méhely L. A magyar fauna Bombinatorjai. II. Hazánk egy új Tritonja: Tr. (Molge) montandoni Blgr. [I. Bombinatoren der ungarischen Fauna. II. Ein neuer Triton: Tr. (Molge) montandoni Blgr./ unseres Landes.]. Mathem Természettud Közl 1892;24:553–74.

[14] Covaciu-Marcov SD, Ferenti S, Bogdan H, Groza MI, Bata ZS. On the hybrid zone between *Bombina bombina* and *Bombina variegata* in Livada Forest, north-western Romania. Biharean Biol 2009;3:5–12.

[15] Szymura JM. Genetic differentiation between hybridizing species *Bombina bombina* and *Bombina variegata* (Salientia, Discoglossidae) in Poland*. Amphibia-Reptilia 1983;4:137–45. https://doi.org/10.1163/156853883X00058.

[16] Gollmann G. *Bombina bombina* and *Bombina variegata* in the Mátra mountains (Hungary): New data on distribution and hybridization (Amphibia, Anura, Discoglossidae). Amphibia-Reptilia 1987;8:213–24. https://doi.org/10.1163/156853887X00252.

[17] Hofman S, Spolsky C, Uzzel T, Cogalniceanu D, Babik W, Szymura JM. Phylogeography of the fire-bellied toads Bombina: independent Pleistocene histories inferred from mitochondrial genomes. Mol Ecol 2007;16:2301–16. https://doi.org/10.1111/j.1365-294X.2007.03309.x.

[18] Fijarczyk A, Nadachowska K, Hofman S, Litvinchuk SN, Babik W, Stuglik M, et al. Nuclear and mitochondrial phylogeography of the European fire-bellied toads *Bombina bombina* and *Bombina variegata* supports their independent histories. Mol Ecol 2011;20:3381–98. https://doi.org/10.1111/j.1365-294X.2011.05175.x.

[19] Vines TH, Köhler SC, Thiel M, Ghira I, Sands TR, MacCallum CJ, et al. The maintenance of reproductive isolation in a mosaic hybrid zone between the fire-bellied toads *Bombina bombina* and *B. variegata*. Evolution 2003;57:1876–88.

[20] Nürnberger B, Barton NH, Kruuk LEB, Vines TH. Mating patterns in a hybrid zone of fire-bellied toads (Bombina): inferences from adult and full-sib genotypes. Heredity (Edinb) 2005;94:247–57. https://doi.org/10.1038/sj.hdy.6800607.

[21] MacCallum CJ, Nürnberger B, Barton NH, Szymura JM. Habitat preference in the bombina hybrid zone in croatia. Evolution (N Y) 1998;52:227–39. https://doi.org/10.1111/j.1558-5646.1998.tb05156.x.

[22] Yanchukov A, Hofman S, Szymura JM, Mezhzherin S V, Morozov-Leonov SY, Barton NH, et al. Hybridization of the *Bombina* and *B. Variegata* (Anura, Discoglossidae) at a sharp ecotone in Westerb Ukraine: comparisons across transects and over time. Evolution (N Y) 2006;60:583–600.

[23] Arnold HR. Atlas of amphibians and reptiles in Britain. Muséum national d’histoire naturelle; 1995.

[24] Harrison RG (Richard G, Biology IC of S and E. Hybrid zones and the evolutionary process. Coll Park Md 1993:364.

[25] Arntzen JW. Some Hypotheses on Postglacial Migrations of the Fire-Bellied Toad, *Bombina bombina* (Linnaeus) and the Yellow-Bellied Toad, *Bombina variegata* (Linnaeus). J Biogeogr 1978;5:339. https://doi.org/10.2307/3038027.

[26] Indykiewicz P. Urban fauna : studies of animal biology, ecology and conservation in European cities. Uniwersytet Technologiczno-Przyrodniczy; 2011.

[27] Szabó I. Adatok a Börzsöny-hegység herpetofaunájához (Data to the herpetofauna of the Börzsöny Mountains). VertebrHung 1960;2:199–216.

[28] Urban P. Poznámky k výskytu oboj ž ivelníkov (Lissamphibia) vybraných antropogénnych vodných nádr ž í Krupinskej planiny 2009.

[29] Sambrook J, Russell DW (David W. Molecular cloning : a laboratory manual. Cold Spring Harbor Laboratory Press; 2001.

[30] Corpet F. Multiple sequence alignment with hierarchical clustering. Nucleic Acids Res 1988;16:10881–90.

[31] Vincze T, Posfai J, Roberts RJ. NEBcutter: A program to cleave DNA with restriction enzymes. Nucleic Acids Res 2003;31:3688–91.

[32] Neff MM, Turk E, Kalishman M. Web-based primer design for single nucleotide polymorphism analysis. Trends Genet 2002;18:613–5.

[33] Pabijan M, Spolsky C, Uzzell T, Szymura JM. Comparative Analysis of Mitochondrial Genomes in Bombina (Anura; Bombinatoridae). J Mol Evol 2008;67:246–56. https://doi.org/10.1007/s00239-008-9123-3.

[34] Wang W, Li J, Wang H, Ran Y, Wu H. Genetic evidence reveals male-biased dispersal of the Omei tree frog. J Zool 2020;310:201–9. https://doi.org/10.1111/jzo.12749.

[35] Hettyey A, Vági B, Kovács T, Ujszegi J, Katona P, Szederkényi M, et al. Reproductive interference between *Rana dalmatina* and *Rana temporaria* affects reproductive success in natural populations. Oecologia 2014;176:457–64. https://doi.org/10.1007/s00442-014-3046-z.

[36] Vági B, Hettyey A. Intraspecific and interspecific competition for mates: *Rana temporaria* males are effective satyrs of *Rana dalmatina* females. Behav Ecol Sociobiol 2016;70:1477–84. https://doi.org/10.1007/s00265-016-2156-5.

[37] Mccarty JP. Ecological Consequences of Recent Climate Change. Conserv Biol 2001;15:320–31.

[38] Walther GR. Community and ecosystem responses to recent climate change. Philos Trans R Soc B Biol Sci 2010;365:2019–24. https://doi.org/10.1098/rstb.2010.0021.

[39] Lindner M, Maroschek M, Netherer S, Kremer A, Barbati A, Garcia-Gonzalo J, et al. Climate change impacts, adaptive capacity, and vulnerability of European forest ecosystems. For Ecol Manage 2010;259:698–709. https://doi.org/10.1016/j.foreco.2009.09.023.

[40] Bellard C, Bertelsmeier C, Leadley P, Thuiller W, Courchamp F. Impacts of climate change on the future of biodiversity. Ecol Lett 2012;15:365–77. https://doi.org/10.1111/j.1461-0248.2011.01736.x.

[41] Carey C, Alexander MA. Climate change and amphibian declines: is there a link? Divers Distrib 2003;9:111–21.

[42] Lips KR, Diffendorfer J, Mendelson JR, Sears MW. Riding the Wave: Reconciling the Roles of Disease and Climate Change in Amphibian Declines. PLoS Biol 2008;6:e72. https://doi.org/10.1371/journal.pbio.0060072.

[43] Blaustein AR, Walls SC, Bancroft BA, Lawler JJ, Searle CL, Gervasi SS. Direct and Indirect Effects of Climate Change on Amphibian Populations. Diversity 2010;2:281–313. https://doi.org/10.3390/d2020281.

[44] Araú Jo MB, Thuiller W, Pearson RG. Climate warming and the decline of amphibians and reptiles in Europe. J Biogeogr 2006;33:1712–1728. https://doi.org/10.1111/j.1365-2699.2006.01482.x.

[45] Pupina A PM. Fire-bellied toads *Bombina bombina* L. (Anura: Bombinatoridae) populations expansion in 2010 in Latvia. Atti IX Congr Naz Della Soc Herpetol Ital Bari-Conversano 2013:343–346.

[46] Dolgener N, Freudenberger L, Schluck M, Schneeweiss N, Ibisch PL, Tiedemann R. Environmental niche factor analysis (ENFA) relates environmental parameters to abundance and genetic diversity in an endangered amphibian, the fire-bellied-toad (*Bombina bombina*). Conserv Genet 2014;15:11–21. https://doi.org/10.1007/s10592-013-0517-4.

[47] Barandun J, Reyer H-U. Reproductive Ecology of Bombina variegata: Habitat Use. Copeia 1998;1998:497–500. https://doi.org/10.2307/1447450.

[48] Cayuela H, Valenzuela-Sánchez A, Teulier L, Martínez-Solano Í, Léna J-P, Merilä J, et al. Determinants and Consequences of Dispersal in Vertebrates with Complex Life Cycles: A Review of Pond-Breeding Amphibians. Q Rev Biol 2020;95:1–36. https://doi.org/10.1086/707862.

[49] Hartel T, von Wehrden H. Farmed Areas Predict the Distribution of Amphibian Ponds in a Traditional Rural Landscape. PLoS One 2013;8:e63649.

